# High resolution structure of the membrane embedded skeletal muscle ryanodine receptor

**DOI:** 10.1101/2021.03.09.434632

**Authors:** Zephan Melville, Kookjoo Kim, Oliver B. Clarke, Andrew R. Marks

## Abstract

The type 1 ryanodine receptor (RyR1)/calcium release channel on the sarcoplasmic reticulum (SR) is required for skeletal muscle excitation-contraction coupling and is the largest known ion channel, comprised of four 565 kDa protomers. Cryogenic electron microscopy (cryoEM) studies of the RyR have primarily used detergent to solubilize the channel, though a recent study resolved the structure with limited resolution in nanodiscs^1^. In the present study we have used cryoEM to solve high-resolution structures of the channel in liposomes using a gel-filtration approach with on-column detergent removal to form liposomes and incorporate the channel simultaneously, a method that improved the incorporation rate by more than 20-fold compared to a dialysis-based approach. In conjunction with new direct-detection cameras, this allowed us to resolve the structure of the channel in the closed and open states at 3.36 and 3.98 Å, respectively. This method offers validation for detergent-based structures of the RyR and lays the groundwork for studies utilizing an electrochemical gradient mimicking the native environment, such as that of the SR, where Ca^2+^ concentrations are millimolar in the lumen and nanomolar in the cytosol of the cell at rest.

## Introduction

Located on the sarco/endoplasmic reticulum (SR/ER) membrane, the two megadalton ryanodine receptor (RyR) is the largest known ion channel^2,3^. In skeletal muscle, RyR1 is required for SR Ca^2+^ release during excitation-contraction coupling (EC coupling). There are three RyR isoforms, RyR1 is the primary isoform in skeletal muscle while RyR2 is the cardiac isoform, though both are found in numerous tissues including neurons^4-6^. RyR3 is widely expressed but at significantly lower levels^7^. In skeletal muscle, RyR1 is thought to be activated by direct interaction with the dihydropyridine receptor (DHPR)^8,9^, whereas RyR2 is activated by Ca^2+^ in a process termed calcium-incuded calcium release (CICR)^10,11^. In short, Ca^2+^ influx via the DHPR triggers the release of Ca^2+^ via RyR2, which in turn creates a high local concentration of Ca^2+^ that activates neighboring RyR2 tetramers. RyR channels are tightly packed in checkerboard arrays on the SR and exhibit cooperative activation/deactivation through the process termed coupled gating^12-15^. Leaky RyR channels are associated with numerous diseases and disorders including muscular dystrophy^16^, heart failure^17^, cardiac arrhythmias^18^, diabetes^19^, Huntington’s Disease^20-22^, and Alzheimer’s^2,3,23,24^.

The high resolution structures of individual, detergent solubilized RyR1^25-28^ and RyR2^29^ channels have been solved using cryogenic electron microscopy (cryoEM)^15^; however, solving the structure of proteins in native or near-native membranes remains challenging. Most cryoEM structures of membrane proteins use detergent micelles, amphipols, or nanodiscs^30-34^. Liposomes, along with linear and circularized nanodiscs, allow for membrane protein incorporation while polymers such as DIBMA and SMA allow for native membrane extractions into so called native nanodiscs^35-38^; however, critically, liposomes are the only method that allows for the creation of asymmetric environments, such as that of the SR. In the lumen of the SR, millimolar concentrations of Ca^2+^ are stored and maintained by the sarco/endoplasmic reticulum Ca^2+^- ATPase (SERCA) and calsequestrin, while the cytosol of the cell remains at approximately 150 nM Ca^2+^ at rest and low micromolar following Ca^2+^ release through the RyR. Liposomes form spontaneously as detergent is removed from solution during dialysis, or through the use of Bio-beads that adsorb detergent molecules. Lige Tonggu & Liguo Wang have shown that liposomes can also be formed via gel filtration^39^. This method allows for significantly greater control over the size of the liposomes formed compared to dialysis^1^ and allowed for the successful incorporation of the large conductance calcium-activated potassium (BK) channel into liposomes^40^.

Given the critical role of the RyR in normal and pathologic physiology, it is important to understand the structure and dynamics of the channel in the native context and to compare these structures to detergent solubilized channels. To this end, the structure and dynamics of proteoliposomal RyR1 was found to match that of detergent, offering validation for the detergent-resolved structures. As with detergent-resolved structures of the RyR, the dynamics of the cytosolic shell undergoes dramatic conformation changes in both the open and closed states of RyR1 channels. These movements are separate from the opening of the channel, marked by the dilation of the pore; however, these changes are difficult to understand in the context of an array of channels, each of which is connected to its neighbors at the four corners of the cytosolic domain^41-43^.

In a truly native context, these dynamics should be muted, but it seems a fully-native approach may be necessary to capture these coupled channels. While we were unable to observe any such coupled channels in liposomes, we have successfully resolved the structure of RyR1 in liposomes in both the closed and open states to 3.36 and 3.98 Å, respectively, through a simple adaption of the gel filtration approach of liposome formation and incorporation. In doing so, we have established a framework for studies more closely mimicking the native environment.

## Results

Our approach, detailed in Methods, is based on the method of Tonggu & Wang^39^ and resulted in small, consistent liposomes compared to forming liposomes by dialysis, which resulted in a wide range of liposome sizes (**Figure 1A**). Critically, this method also offered a greater than 20-fold increase in the RyR1 incorporation rate based on the average number of particles per micrograph with approximately 8 channels per micrograph compared to 1 particle per 3 micrographs observed in the dialysis-based approach. Unfortunately, while both methods of liposome formation required an additional clearing step, liposomes formed via gel filtration showed only minor improvement from passage over an ion-exchange column. While many particles of unincorporated RyR1 were still present, these were discarded during 2d classification by increasing the box size to ensure the membrane would be clear and present as well as increasing the number of classes to weed out unincorporated RyR1. Despite the lack of top and bottom views of the channel, 360° of side views were present in addition to oblique views (**Figure S2**), allowing for a full 3d reconstruction of the channel in a lipid membrane, showing the potential of this method to achieve high-resolution cryoEM structures of proteins in membranes.

**Figure 1.**
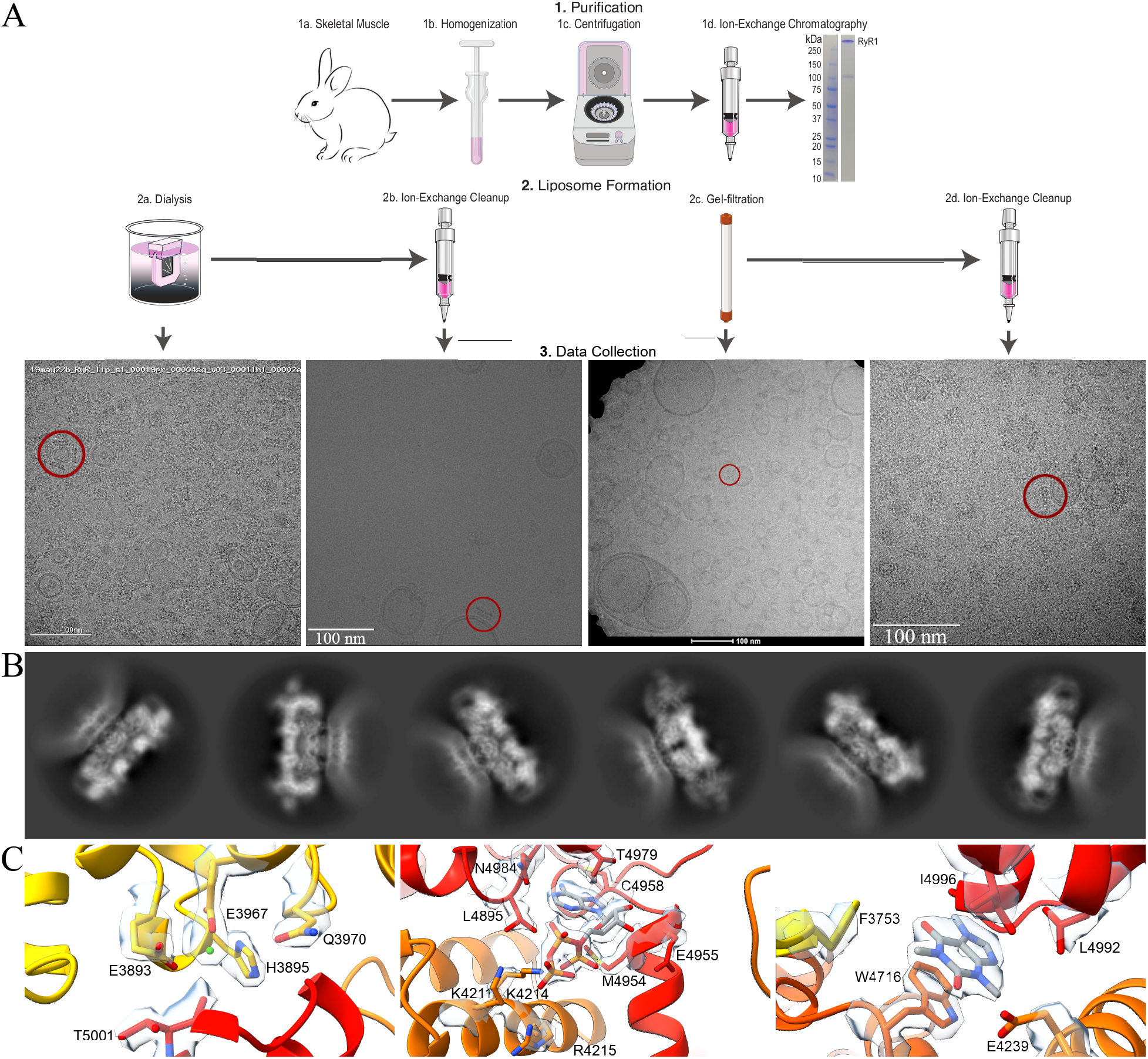
Workflow diagram for the preparation of proteoliposomal RyR1. ***A***. RyR1 was purified from rabbit skeletal muscle using ion-exchange chromatography prior to liposome formation and incorporation by dialysis or gel filtration. For both methods, a second round of ion-exchange chromatography was employed to eliminate unincorporated RyR1 and empty liposomes. An embedded channel is highlighted in representative micrographs. ***B***. 2D classes of proteoliposomal RyR1 showing the flattening of the lipid membrane around the TM domain of the channel. ***C***. Representative density for the Ca^2+^, ATP, and caffeine binding sites of RyR1 in the closed state of proteoliposomal RyR1, resolved to 3.36 Å.

The structure of proteoliposomal RyR1 matches the detergent-resolved structure^26^ with the exception of the hydrophobic-gate residue, I4937. In liposomes, this pore helix rotates such that I4937 is no longer within the pore in the open state of the channel. As a result, the radius of the pore residue cannot not be measured at I4937 (**Figure 2**)^44^, though by estimation it remains at approximately 4 Å, which is consistent with the 4 Å radius or 8 Å diameter observed in detergent. The lumen of RyR1 at the base of the channel varies between liposomes and detergent but this region consists of unstructured loops so dynamic behavior is to be expected. Analysis of the calstabin binding site, along with the binding sites for calcium, ATP, and caffeine show no significant changes between detergent solubilized and liposome embedded RyR1 channels for both the closed and open states, however the improved resolution in the closed state allows for unambiguous assignment of the orientation of caffeine within the binding site (**Figure 1C**).

**Figure 2.**
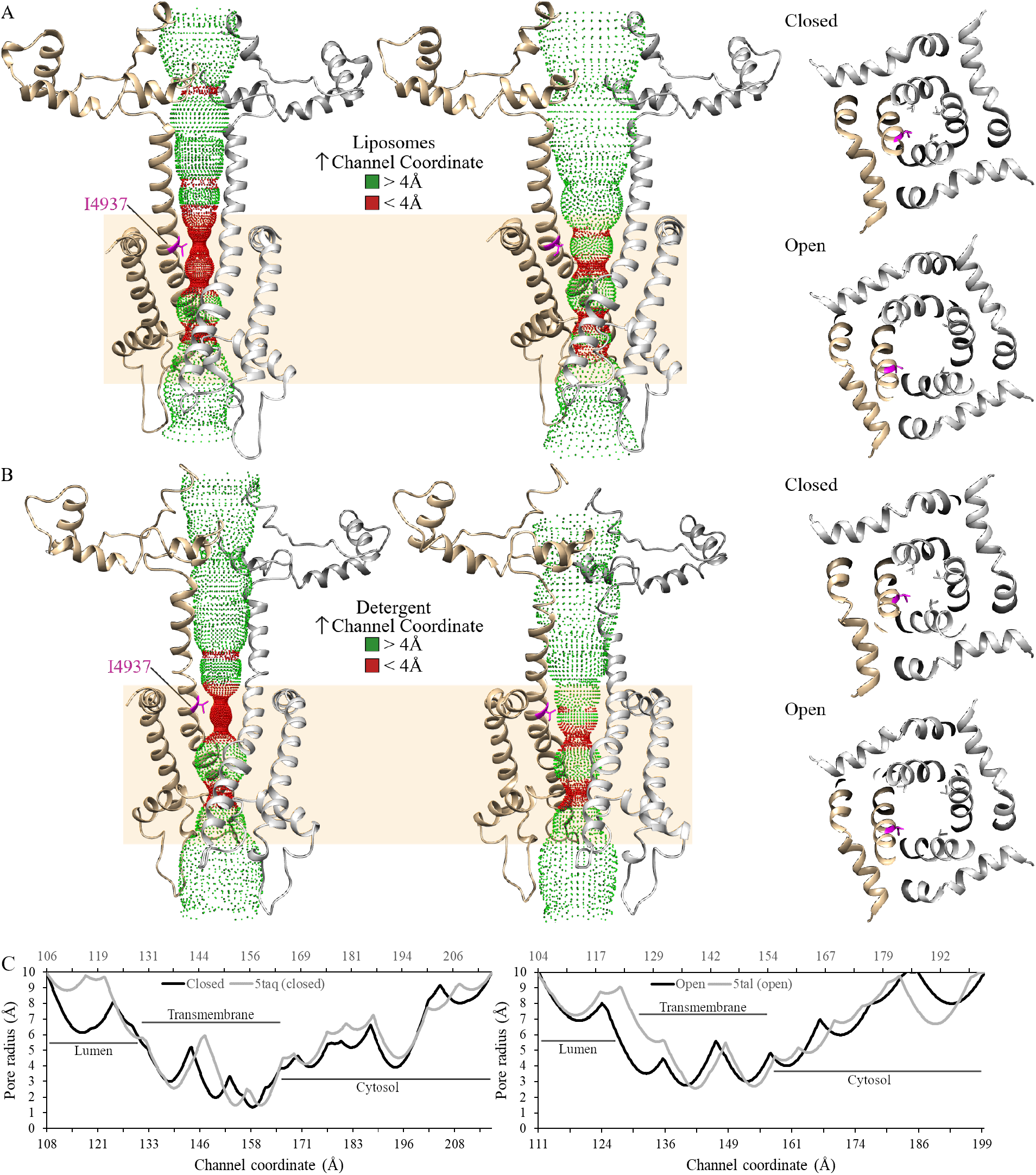
Accessible inner surface of the RyR1 pore. The transmembrane pore (residues 4820-5037) is depicted as a ribbon diagram of two protomers with the hydrophobic gate residue, I4937, in magenta. The dotted representation of the accessible inner surface of the channel was calculated using HOLE and is colored green where the radius exceeds 4 Å and red when the radius is less than 4 Å. The transmembrane domain of RyR1 is highlighted in tan. *A*. Side and cytoplasmic views of the inner surface of the pore of proteoliposomal RyR1 in the closed and open states. *B*. Side and cytoplasmic views for the RyR1 in detergent in the closed (5taq) and open (5tal) state. *C*. Graphical representation of the pore radius of the channel in liposomes (black) and detergent (gray) in the closed and open states. The channel coordinates follow from the lumen to the cytosolic side of the RyR1 with the liposomal structure on the primary x-axis and the detergent structures on the secondary x-axis.

The juxtamembrane helices at the top of the TM interface in each protomer make several contacts with the lipid membrane, however the lipid membrane is too dynamic to clearly identify these interactions. These helices are likely responsible for the flattening of the lipid membrane around the TM domain of the RyR that was consistently present in 2D classes (**Figure 1B**). Likewise, the dynamics of the cytosolic shell of the liposome embedded channel conform to what has previously been observed. The detergent solubilized RyR1 structure undergoes dramatic conformation changes that are separate from channel opening, in both the closed and open states (**Supplementary Movie 1&2**)^25,26^. The cytosolic shell of RyR1 rises and falls analogous to flapping wings. This movement is matched in the liposome embedded channels (**Supplementary Movie 3&4**), though these movements are likely to be exaggerated in the absence of an array of channels.

## Discussion

The present study provides refined methodology that can be applied to other membrane proteins including receptors and channels. Additional cleanup steps may be required to eliminate empty liposomes and denatured protein.Since functional studies of RyR1 are often performed using planar lipid bilayers^45^, solving the structure in a similar membrane establishes that these channels are likely in physiologically relevant conformations. The ability to study the structure of the channel in proteoliposomes now provides the opportunity to resolve structures of membrane proteins under markedly more physiological conditions. In the case of RyR1, the Ca^2+^ concentration in the lumen of the SR is millimolar, but the cytosolic shell is exposed to nanomolar Ca^2+^ at rest. Ca^2+^ within the SR is not simply free Ca^2+^, but is significantly bound to Ca^2+^ binding proteins including calsequestrin and going forward it may be possible to incorporate calsequestrin into the lumen of proteoliposomes for structural studies.

One limitation of the present study is that the structure of liposome embedded RyR1 channels remains limited to single RyR1 channels, while native RyR1 exists in an array of channels in a checkerboard pattern on the terminal cisternae of the SR membrane^41-43^. Although as many as three molecules of RyR1 were sometimes present in a single liposome, no intermolecular interactions were observed^46-48^. Curiously, multiple channels were only observed in small liposomes (<100 nm diameter); however, no more than a single molecule of RyR1 was observed in liposomes >100 nm. Instead, larger liposomes were more likely to be empty. Thus, a structure of RyR1 in native SR membranes will be necessary to observe RyR1 arrays and the native intermolecular dynamics. In addition to validation of existing structures of detergent solubilized RyR1, the present study provides a road map for studies of RyR1 structures in native membranes at high resolution using cryoEM^49^, including studies mimicking the electrochemical gradient between the cytoplasm of the cell and the lumen of the SR.

## Methods

The workflow for solving the high-resolution structure of RyR1 embedded in liposomes is outlined in Figure S1.

### RyR1 purification

All purification steps were performed on ice unless otherwise stated. RyR1 was purified from rabbit skeletal muscle as described previously with modifications^50^. Rabbit back and thigh skeletal muscle, purchased from BioIVT, was snap frozen in liquid nitrogen immediately following euthanasia prior to storage at -80°C. 20 g of rabbit skeletal muscle was homogenized in a Waring blender and lysed in buffer A [10 mM tris maleate pH 6.8, 1 mM EGTA, 1 mM benzamidine hydrochloride, 0.5 mM 4-(2- aminoethyl)benzenesulfonyl fluoride hydrochloride (AEBSF)]. The resulting suspension was pelleted by centrifugation for ten minutes at 11,000 X g. The supernatant was filtered through cheesecloth to remove debris and the membranes were then pelleted by centrifugation for thirty minutes at 36,000 X g. The membranes were then solubilized in buffer B [10 mM HEPES pH 7.4, 0.8 M NaCl, 1% CHAPS, 0.1% phosphatidylcholine, 1 mM EGTA, 2 mM DTT, 0.5 mM AEBSF, 1 mM benzamidine hydrochloride, 1 protease inhibitor tablet (Pierce)] prior to homogenization using a glass tissue grinder (Kontes). Homogenization was repeated following the addition of buffer C (buffer B with no NaCl) at a 1:1 mixture with buffer B. The resulting homogenate was subjected to centrifugation for thirty minutes at 100k X g. The supernatant was then vacuum filtered and loaded at 1 mL/min onto a HiTrap Q HP column (5 mL, GE Healthcare Life Sciences) pre-equilibrated with buffer D [10 mM HEPES pH 7.4, 400 mM NaCl, 1.0% CHAPS, 1 mM EGTA, 0.5 mM tris(2-carboxyethyl)phosphine (TCEP), 0.5 mM AEBSF, 1 mM benzamidine hydrochloride, 0.01% 1,2-dioleoyl-sn-glycero-3-phosphocholine (DOPC, Avanti)]. DOPC (25 mg/mL), dissolved in chloroform, was evaporated under nitrogen gas and resuspended in buffer D. Contaminating proteins were washed away with six column volumes (CV) of buffer D prior to elution of RyR1 with a linear gradient from 480 to 550 mM NaCl using buffers D and E (buffer D with 600 mM NaCl).

### Liposome formation using dialysis and initial screening

Purified RyR1 (1 mg/mL) was incubated at a ratio of 1:2,000 with a 5:3 ratio of phosphatidylethanolamine and phosphatidylcholine and dialyzed into buffer F [10 mM HEPES pH 7.4, 400 mM NaCl, 1 mM EGTA, 0.5 mM TCEP] for 48 hours replacing the buffer with fresh buffer twice per day. Initial screening was performed on a Tecnai F20 (FEI) microscope at Columbia University. Liposomes formed by dialysis were found to have widely variable sizes and even with passage through an extruder, many liposomes were not unilammelar.

### Liposome formation using gel filtration

Purified RyR1 was concentrated to <0.5 mL and incubated with 1:1,000 5:3 phosphatidylethanolamine and phosphatidylcholine in 10% CHAPS for 10 minutes prior to loading onto a hand-packed G50 column pre-equilibrated with buffer F. Liposome formation and RyR1 incorporation occurred on the column and proteoliposomal RyR1 was eluted at 0.20-0.25 mL/min in buffer F.

### Cleanup

Proteoliposomal RyR1 was cleared via passage over a second HiTrap Q column, washed at 425 mM NaCl then eluted at 600 mM NaCl. Proteoliposomal RyR1 was concentrated to 2.5 mg/mL, determined by spectroscopy using a NanoDrop 1000 (ThermoFisher, 1 abs @ 280 nm = 1 mg/mL).

### Grid preparation

UltrAuFoil holey gold grids (Quantifoil R 0.6/1.0, Au 300) were plasma cleaned for thirty seconds with H_2_ and O_2_ (Gatan). Prior to setting grids, proteoliposomal RyR1 was incubated for 10 min with 2 mM NaATP, 5 mM caffeine, and 30 µM Ca^2+^_free_ (MaxChelator)^51^. These conditions were chosen to ensure both the closed and open states would be visible. 3.0 µL of 2.5 mg/mL proteoliposomal RyR1 was applied to each grid. Grids were then blotted for 7.5 sec at blot force 3, with a wait time of thirty seconds prior to vitrification by plunge freezing into liquid ethane chilled with liquid nitrogen^52,53^ with a Vitrobot Mark IV (ThermoFisher) operated at 4°C with 100% relative humidity.

### Data Collection

Grids prepared with liposomes formed by gel filtration were screened at City University of New York (CUNY) using a 120-kV G2 Spirit Twin microscope (FEI Tecnai). Microscope operations and data collection were carried out using the SerialEM software^54^. High resolution data collection was performed at Columbia University on a Titan Krios 300-kV (ThermoFisher) microscope equipped with an energy filter (slit width 20 eV) and a K3 direct electron detector (Gatan). Data were collected using Leginon^55^ and at a nominal magnification of 105,000X in electron counting mode, corresponding to a calibrated pixel size of 0.828 Å. The electron dose rate was set to 16 e^-^/pixel/sec with 2.5 second exposures, for a total dose of 58.34 e/A^2^. These grids showed small, consistent liposomes, similar to those reported by by Tonggu & Wang^39^ and significantly greater incorporation rate of RyR1; however, empty liposomes and unincorporated and aggregated RyR1 also remained.

### Data Processing

CryoEM data processing was performed using cryoSPARC^56-60^ with the exception of 3D classes, which were performed in Relion 3.1 to ensure accurate comparisons of the dynamics between liposome embedded channels and the previously published detergent-resolved channels^26,61^. Image stacks were aligned using Patch motion correction and defocus value estimation by Patch CTF estimation. Initial particle picking was performed manually for >100 particles to create templates prior to template-based picking. This involved increasing the box size to ensure a portion of the lipid membrane would be included. Two million particles were initially picked from 11,187 micrographs and these were subjected to several iterations of 2D classification in cryoSPARC with 200 classes each to separate proteoliposomal RyR1 from empty liposomes and free RyR1. Classes comprised of 175,000 particles of unincorporated RyR1 were eliminated during iterative rounds of 2D classification, leaving 85,000 liposome-embedded particles. This represents a greater than 20-fold increase in RyR1 incorporation rate over the dialysis-based method which resulted in 2,600 particles from over 7,000 micrographs. The particles from the highest-resolution classes were pooled for ab initio 3D reconstruction with a single class followed by homogenous refinement with per-particle defocus refinement and C4 symmetry imposed. 3D variability analysis revealed the presence of the open state and heterogenous refinement was performed in order to separate the two states, with 63% of particles (53,882) in the closed state and 37% in the open state (31,599).

Symmetry expansion and local refinement, were performed using cryoSPARC to improve local resolution. Local refinement was performed using three separate masks. The first mask was composed of the N-terminal domain, the SPRY domains, the RY1&2 domain, and calstabin. The second mask surrounded the bridging solenoid, and the third mask surrounded the RyR1 pore. The resulting maps were combined in Chimera^62^ to generate a composite map prior to calibration of the pixel size using the crystal structure of the N-terminal domain (2XOA)^63^. Model building was performed in Coot^64^ with refinement in Phenix^65,66^ and figures were generated using Chimera^62^, ChimeraX^67^, and PyMol^68^. The pore aperture of RyR1 was calculated using HOLE^44^. Movies of the dynamics of the channel were made in ChimeraX using maps generated by 3D classes for liposomes and the corresponding maps from 3D Classes in detergent (5TAN, 5TAM, 5TA3, 5T9V). Each map was filtered to 5 Å and aligned at the pore. CryoEM statistics are summarized in **Figure S2** and **Table S1**.

## Supporting information

Supplemental movie 1 RyR1 detergent closed

Supplemental movie 2 RyR1 detergent open

Supplemental movie 3 Proteoliposomal RyR1 closed

Supplemental movie 4 Proteoliposomal RyR1 open

## Acknowledgements

These studies were supported by NIH grants R01HL145473, R01DK118240, R01HL142903, R01HL140934, R01AR070194 and T32 HL120826 (to A.R.M.).

## Author contributions

Z.M. conducted experiments, analyzed data, and wrote the manuscript; K.K. conducted experiments and analyzed data; A.R.M and O.B.C. analyzed data and edited the manuscript.

## Figure Legends

**Figure S1.**
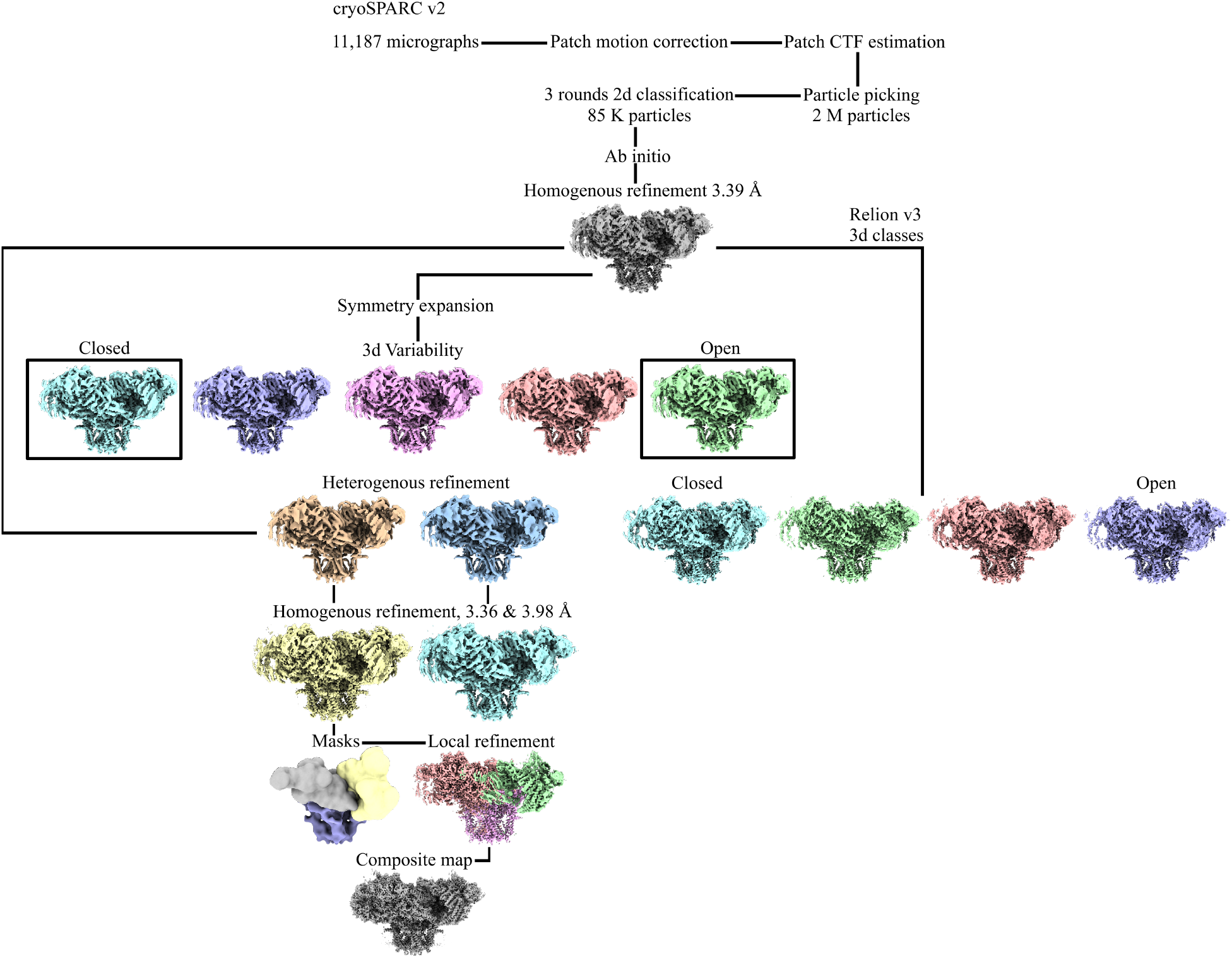
Data processing workflow for proteoliposomal RyR1 in cryoSPARC and Relion.

**Figure S2.**
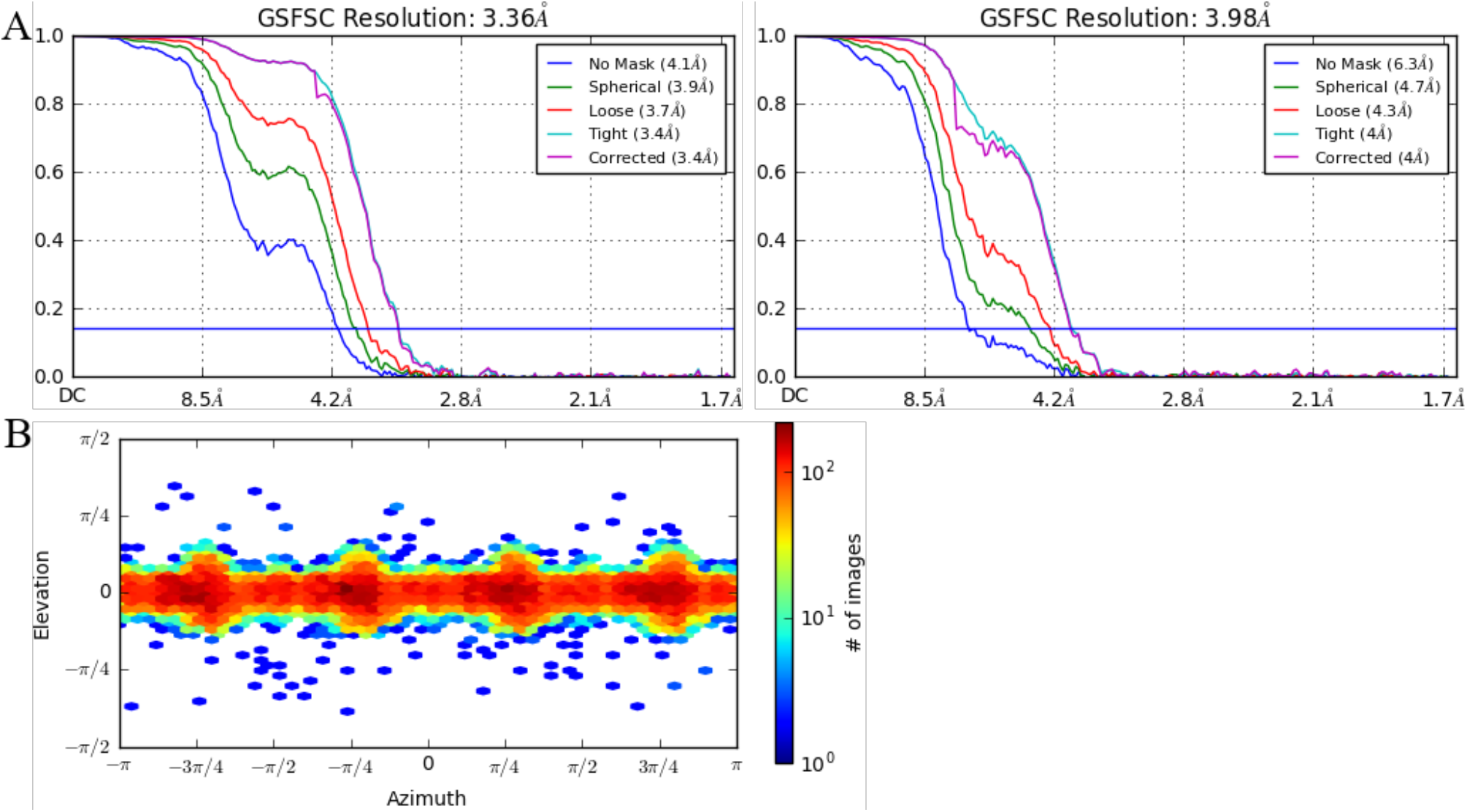
*A*. GSFSC curves of proteoliposomal RyR1 in the closed (left) and open (right) states. *B*. Viewing angle distributions of proteoliposomal RyR1.

**Table S1.**
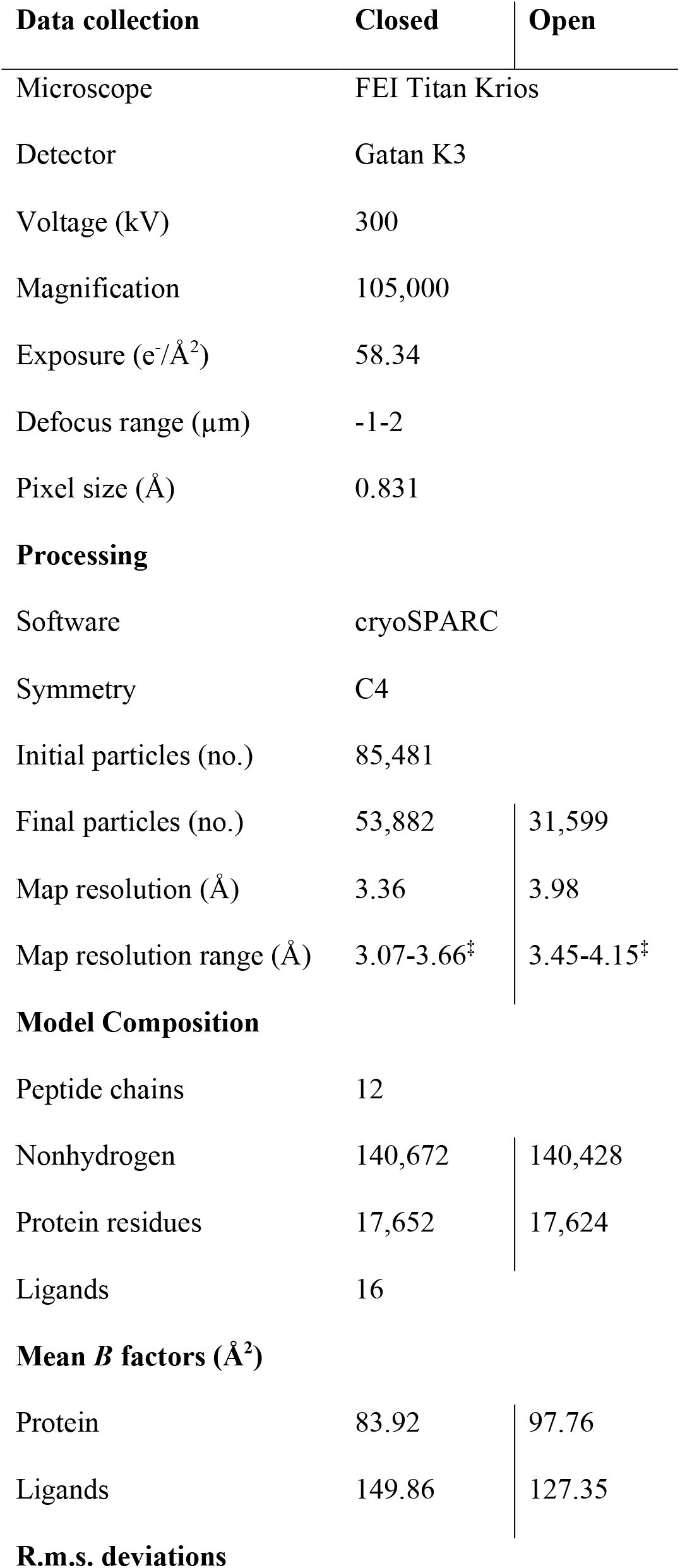
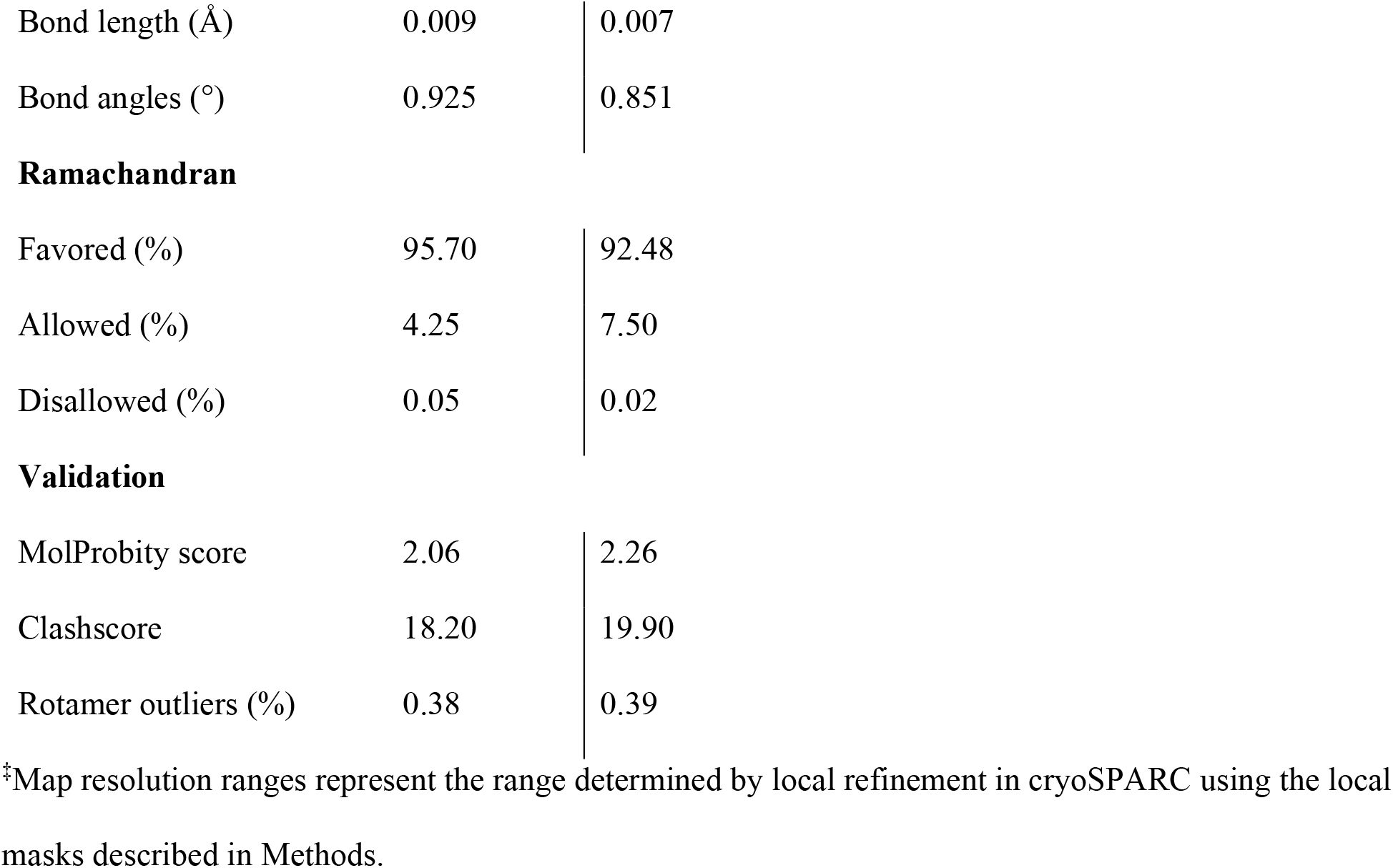

| Data collection | Closed | Open |
| --- | --- | --- |
| Microscope | FEI Titan Krios |  |
| Detector | Gatan K3 |  |
| Voltage (kV) | 300 |  |
| Magnification | 105,000 |  |
| Exposure (e<sup>-</sup>/Å<sup>2</sup>) | 58.34 |  |
| Defocus range (μm) | -1-2 |  |
| Pixel size (Å) | 0.831 |  |
| Processing |  |  |
| Software | cryoSPARC |  |
| Symmetry | C4 |  |
| Initial particles (no.) | 85,481 |  |
| Final particles (no.) | 53,882 | 31,599 |
| Map resolution (Å) | 3.36 | 3.98 |
| Map resolution range (Å) | 3.07-3.66<sup>‡</sup> | 3.45-4.15<sup>‡</sup> |
| Model Composition |  |  |
| Peptide chains | 12 |  |
| Nonhydrogen | 140,672 | 140,428 |
| Protein residues | 17,652 | 17,624 |
| Ligands | 16 |  |
| Mean *B* factors (Å<sup>2</sup>) |  |  |
| Protein | 83.92 | 97.76 |
| Ligands | 149.86 | 127.35 |
| R.m.s. deviations |  |  |
| Bond length (Å) | 0.009 | 0.007 |
| Bond angles (°) | 0.925 | 0.851 |
| **Ramachandran** |  |  |
| Favored (%) | 95.70 | 92.48 |
| Allowed (%) | 4.25 | 7.50 |
| Disallowed (%) | 0.05 | 0.02 |
| **Validation** |  |  |
| MolProbity score | 2.06 | 2.26 |
| Clashscore | 18.20 | 19.90 |
| Rotamer outliers (%) | 0.38 | 0.39 |

‡Map resolution ranges represent the range determined by local refinement in cryoSPARC using the local masks described in Methods.

